# Controllability in an islet specific regulatory network identifies the transcriptional factor NFATC4, which regulates Type 2 Diabetes associated genes

**DOI:** 10.1101/226456

**Authors:** Amitabh Sharma, Arda Halu, Julius L. Decano, Jörg Menche, Yang-Yu Liu, Rashmi B. Prasad, Joao Fadista, Marc Santolini, Megha Padi, Scott T. Weiss, Marc Vidal, Edwin K. Silverman, Masanori Aikawa, Albert-László Barabási, Leif Groop, Joseph Loscalzo

## Abstract

Probing the dynamic control features of biological networks represents a new frontier in capturing the dysregulated pathways in complex diseases. Here, using patient samples obtained from a pancreatic islet transplantation program, we constructed a tissue-specific gene regulatory network and used the *control centrality (Cc)* concept to identify the high control centrality (HiCc) pathways, which might serve as key pathobiological pathways for Type 2 Diabetes (T2D). We found that HiCc pathway genes were significantly enriched with modest GWAS *p*-values in the DIAbetes Genetics Replication And Meta-analysis (DIAGRAM) study. We identified variants regulating gene expression (expression quantitative loci, eQTL) of HiCc pathway genes in islet samples. These eQTL genes showed higher levels of differential expression compared to non-eQTL genes in low, medium and high glucose concentrations in rat islets. Among genes with highly significant eQTL evidence, NFATC4 belonged to four HiCc pathways. We asked if the expressions of T2D-associated candidate genes from GWAS and literature are regulated by Nfatc4 in rat islets. Extensive *in vitro* silencing of Nfatc4 in rat islet cells displayed reduced expression of 16, and increased expression of 4 putative downstream T2D genes. Overall, our approach uncovers the mechanistic connection of NFATC4 with downstream targets including a previously unknown one, TCF7L2, and establishes the HiCc pathways’ relationship to T2D.

## Introduction

The pathobiological changes leading to a complex disease are most likely to be influenced by the disease genes that perturb the underlying biological networks in specific tissue types. Recent evidence suggests that these perturbations are not scattered randomly in the interactome; instead, they are localized in specific neighborhoods, or ‘disease modules’ (1,2). In order to identify this disease-specific interactome neighborhood, we previously integrated human islet gene expression data, genetics, and protein interaction data to build a localized map of genes associated with islet cell dysfunction in T2D (3). Recently, we identified an asthma disease module by a connectivity-based model and validated it for functional and pathophysiological relevance to the disease (2). Several tools based on the ‘guilt-by-association principle’ predict potential candidate genes using networks (4–6). Furthermore, inference tools such as ANAT identify the highest-confidence paths between pairs of proteins by viewing the local neighborhood of a set of proteins (7). Other methods such as HotNet2 use the heat diffusion process to analyze a gene’s mutation score and its local topology together to find the subnetworks in cancer (8). Similarly, the NetQTL approach combines eQTL and network flow to identify genes and dysregulated pathways (9).

Despite extensive interest in using topological features to interpret the biological networks in human disease, an important aspect that has been largely overlooked so far is the *controllability* of these subcellular networks. In general, controllability can be achieved by changing the state of a small set of driver nodes that govern the dynamics of the entire network (10). Liu *et al*. proposed an analytical framework to identify the minimum set of driver nodes (MDS) of any complex network, whose time-dependent control can guide the whole network to any desired final state. They found that driver nodes tend to avoid hubs, i.e., highly connected nodes(11). Furthermore, Milenkovic et al. suggested the notion of domination and found dominating sets (DSs) in the undirected protein interaction network(12). Wuchty identified the minimum dominating sets (MDSets) that play a role in the control of the underlying protein interaction network^9^. It was observed that MDSet proteins were enriched with cancer-related and virus-targeted genes (13). We recently showed that the application of network controllability analysis helps in identifying new disease genes and drug targets.(14).

Progress towards a robust network-based controllability approach will ultimately lead to the identification of potential key regulatory nodes that govern network function in health and disease. As a first step in this direction, here, we asked whether the set of genes that are predicted to control a biological directed network would affect the functional pathophenotype. To assess the controllability of the network, we used the *control centrality* (Cc) measure (15), which quantifies the ability of a single node to control an entire directed weighted network (see Methods). Our disease of interest, T2D, is a complex disease and therefore has the potential to lend itself to this approach where controllability in a regulatory network specific to it might reveal new knowledge about the disease. T2D is characterized by insufficient insulin secretion from the *β*-cells of islets in the pancreas(3). We hypothesize that the high control centrality (HiCc) pathways, representing specific gene sets in a T2D-regulatory network in human pancreatic islets, might control other downstream pathways involved in disease manifestation (see Figure 1). To test our hypothesis we construct a pancreatic islet-specific extended gene-regulatory network, and use control centrality to identify HiCc pathways in the KEGG database. We validate the disease relevance of these HiCc pathways using T2D-specific -omics data. Next, we test whether the SNPs located in non-coding regions of HiCc pathway genes in islet samples would influence RNA-seq expression (eQTL). Finally, we perform extensive in vitro silencing experiments on NFATC4, an eQTL-implicated gene found in four HiCc pathways, and probe whether T2D-associated genes from GWAS and literature are downstream targets of NFATC4. Overall, our study provides a unique framework for integrating control principles towards distinguishing pathways and genes that are likely to contribute to T2D pathogenesis.

**Figure 1:**
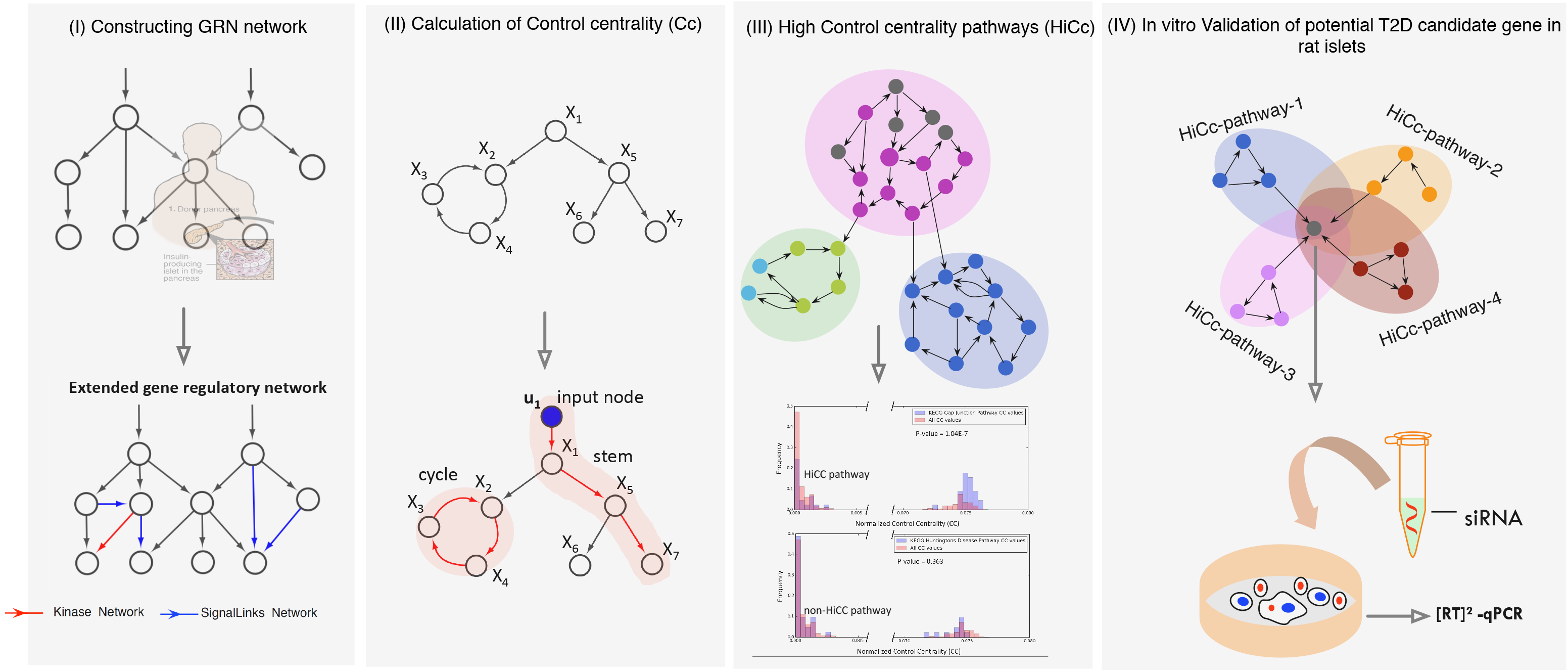
Overview of the approach to identify the key pathways in T2D using control centrality approach. a. Gene expression data: Pancreatic islets from cadaver donors (54 nondiabetic and 9 diabetic) were used to construct the gene regulatory network (GRN) and extended by adding kinase and signaling links. The largest connected component of the extended GRN (EGRN) consists of *N*=3,084 genes and *M*=7,935 edges. b. The *control centrality* measure is used to quantify the relative importance of each gene in EGRN relative to T2D. c. High control centrality (HiCc) pathways are found by comparing the control centrality distribution of genes within the pathway versus the control centrality distribution of all other genes in EGRN. Pathways with a significantly higher control centrality distribution compared to the background are deemed HiCc pathways. For example, the Gap junction pathway emerges as a HiCc pathway, whereas the Huntington’s Disease pathway is found to be a non-HiCc pathway. d. *In vitro* silencing experiments are performed on genes implicated in a large number of HiCc pathways, such as NFATC4, to discover novel mechanistic connections with known T2D genes.

## Results

### Extended Gene regulatory network (EGRN) from human islet cells

We start by building a gene-regulatory network (GRN) using gene expression data from pancreatic islet samples of diabetic and non-diabetic cadaver donors obtained through the Nordic Islet Transplantation Programme (http://owww.nordicislets.org). The GRN consists of differentially expressed genes in diabetic and high glycated hemoglobin (HbA1c) donors, highly varying genes in all donor islets, and established T2D genes from genome-wide association studies (GWAS). Directed edges in the GRN are inferred using a combination of linear regression and prior knowledge from the TRANSFAC database (http://www.gene-regulation.com/). The GRN specific to islet cells consists of 896 genes with 1,164 links between them. We next include most of the signaling events by extending the GRN with the addition of kinase and signaling pathways (see Methods for further details on the construction of the networks). The largest connected component (LCC) of the extended GRN (EGRN) with kinase-substrate and signaling links consists of N=3,084 genes and M=7,935 edges. The average number of neighbors in the network is 5.14. Compared to randomized networks constructed using degree-preserving randomization, the EGRN has a significantly higher average shortest path length <*l*> = 4.65 (z-score=56) (Figure 2a) and a significantly higher clustering coefficient C = 0.055 (z-score=6.86) (Figure 2b). Interestingly, irrespective of the difference in network size and the degree distributions of GRN and EGRN (Supplementary Figure 1a), their normalized control centrality (*c*_c_=*C*_c_/*N*) distributions do not differ significantly (Supplementary Figure 1b). This indicates the robustness of the control centrality measure to changes in the size of the network. Moreover, EGRN’s normalized control centrality is significantly higher compared to randomized networks. The p-values between randomized networks and EGRN using Mann-Whitney U (100 p-values) are between 2.77e-15 and 9.76e-12 (Figure 2c). This suggests that the overall structure of the EGRN is more controllable than its randomized counterparts.

**Figure 2:**
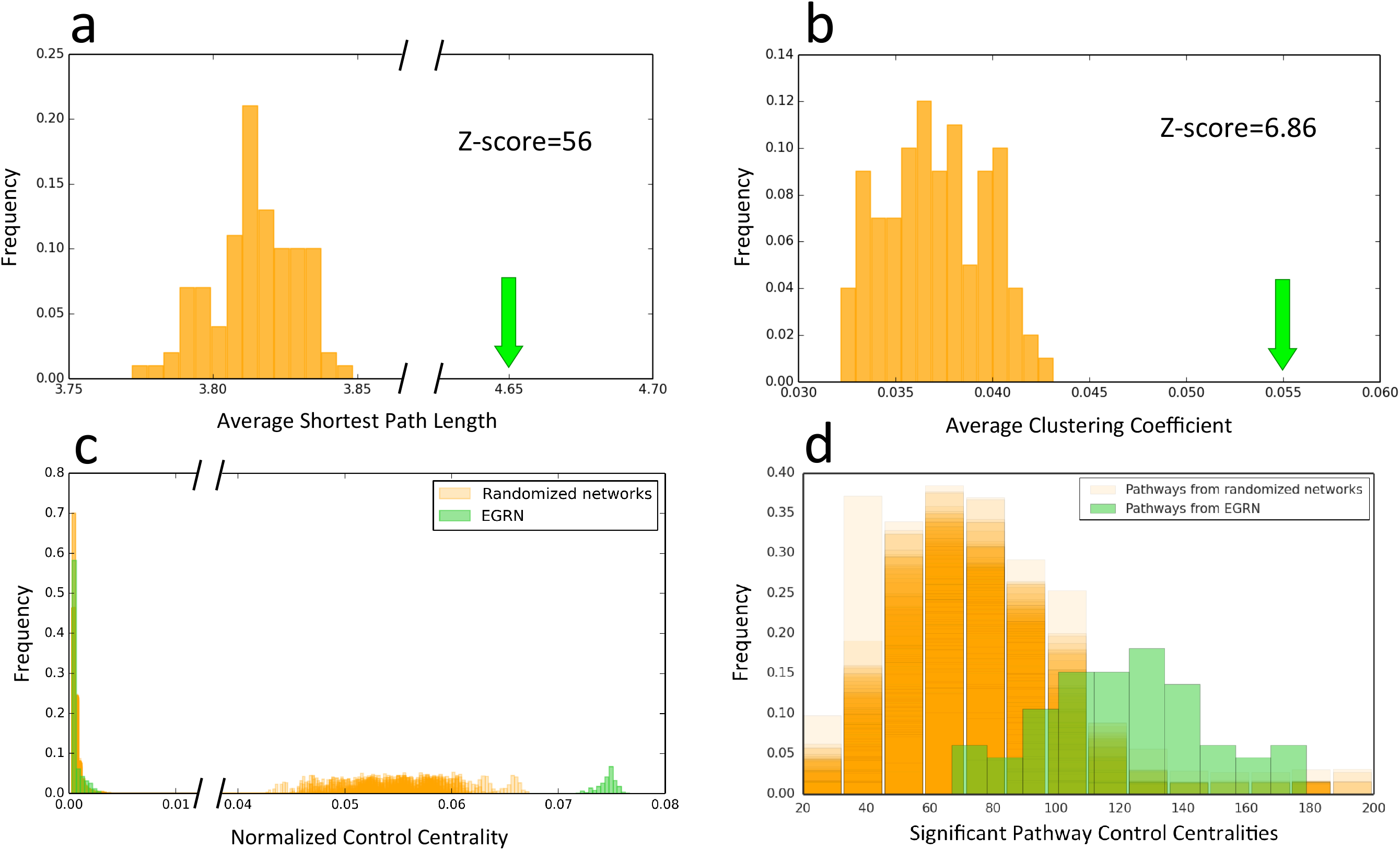
Topological and control centrality-related properties of EGRN. a. The average shortest path length of the EGRN is 4.65, which is significantly higher that those of randomized networks (shown in orange) with a z-score of 56. b. The average clustering coefficient of the EGRN is 0.055, which is significantly higher that those of randomized networks (shown in orange) with a z-score of 6.86. c. The normalized control centrality distribution of the EGRN (shown in green) is significantly higher than those of randomized networks (shown in orange). d. The control centralities of the HiCc pathways derived from the EGRN (shown in green) are significantly higher than those of the HiCc pathways derived from randomized networks.

### Identifying the high control centrality (HiCc) pathways in the EGRN

A complex disease such as T2D is likely to be the result of multiple gene perturbations within pathways in a biological network, where changes in one pathway might trigger alterations in other downstream pathways. Hence, identification of ‘key driver pathways’ in the islet-specific EGRN should give us insights about the molecular processes responsible for the disease. Here, we compared the control centrality distribution of the genes in each pathway in the KEGG database with the control centrality distribution of all other genes in the EGRN, and observed 66 significant HiCc pathways with a *p*-value <0.05 (Mann-Whitney test) (Supplementary Table 2) (for details see Methods). Overall, the genes representing T2D pathways in the EGRN had higher C_c_ values compared to the random distribution (Supplementary Figure 2), indicating that control centrality is able to capture the important pathways associated with T2D in KEGG. To ensure that our observations regarding the role of control centrality on EGRN in teasing out biologically relevant pathway information cannot be reproduced from randomized data, we repeated the control centrality calculations on degree-preserved randomized networks. We found that the control centrality (C_c_) values of the significant pathways are higher on average for EGRN than randomized networks. The average of the means of the Cc distributions for randomized networks was 72.49, whereas the mean of the Cc distribution for the EGRN network was 121.32. The Mann-Whitney U p-values were between 9.54e-24 and 6.78e-11 (Figure 2d). This both proves the utility of using the EGRN in conjunction with control centrality and establishes control centrality as an effective metric for prioritizing pathways.

We next asked whether the genes with the high Cc genes were also hubs., i.e. highly connected genes. We, therefore, compared the degree distributions of: 1) all genes in the EGRN, 2) all genes that are in any of the 66 significant HiCc pathways, and 3) all genes that are in any of the remaining 120 non-significant Cc pathways. Overall, we did not observe any significant differences between the three types of degree distributions as shown in Figure 3a. Thus, both the significant and the nonsignificant pathways based on Cc values contain genes that have similar degree distribution in the EGRN, indicating an absence of bias towards hubs as high Cc genes.

**Figure 3:**
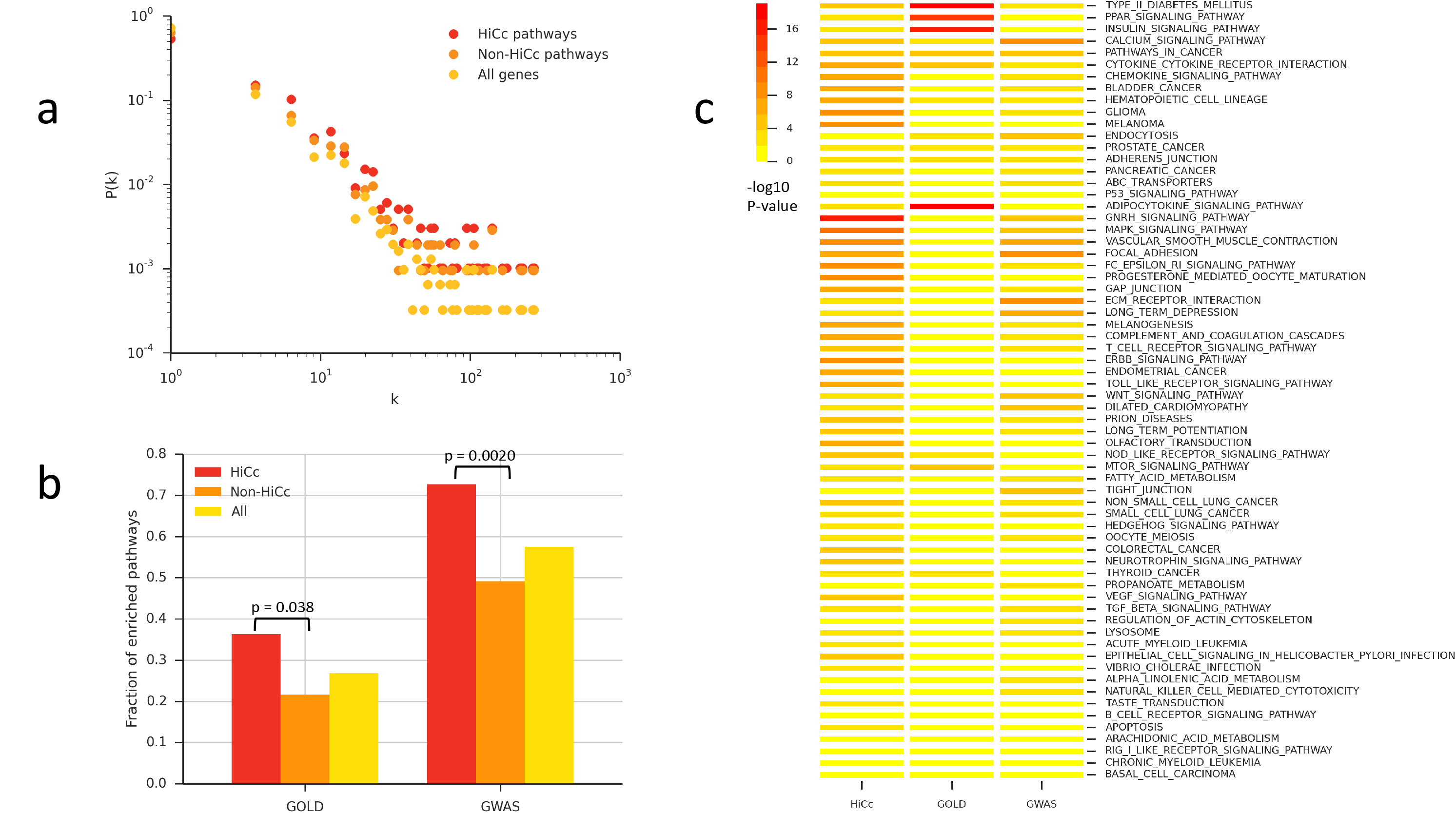
Properties and T2D relevance of high control centrality (HiCc) pathways. a. Degree distributions P(k) of HiCc pathway genes, non-HiCc pathway genes, and all other genes in the EGRN. b. The fraction of enriched pathways in the manually curated GOLD T2D gene data, T2D differential expression (DE) data, and GWAS data, for HiCc pathways, non-HiCc pathways, and all pathways. c. The 66 HiCc pathways and their enrichment in T2D specific data sources.

To test the reliability and performance of the control centrality approach as a means to glean key drivers of T2D, we compared it to other methods that identify the dysregulated subnetworks associated with a specific phenotype. We found that control centrality is comparable to or higher performing than a number of established methods to find the dysregulated pathways, such as HotNet2(8) in capturing “T2D-related” pathways, which are pathways significantly enriched in literature-mined T2D disease genes with experimental evidence from the DISEASES database(16), both on the EGRN and on generic networks (see Supplementary Information for details on comparisons with other methods).

To further assess the performance of control centrality as a network centrality measure, we carried out the T2D diabetes “T2D-related” pathway assessment on other centrality measures applied on the EGRN. We found that control centrality is superior to all of the tested centrality measures in terms of the significance of overlap with T2D-related pathways, i.e. the high control centrality pathways have a higher enrichment of T2D-related pathways than pathways with high centrality according to other centrality measures (Supplementary Table 1). The fact that control centrality outperforms degree centrality also confirms our observation that control centrality, which is not biased towards highly connected nodes, uncovers disease-related information that is independent of the “hubness” of a node.

### T2D relevance of the HiCc pathways in the EGRN

We hypothesized that if HiCc pathways contribute to the control of disease-related processes in T2D, they should be significantly enriched within T2D-specific -omics data. To test this hypothesis, we separated the HiCc pathways, i.e. pathways whose genes have significantly higher Cc values than the rest of the genes in EGRN, from those that are not HiCc. For both groups of pathways, as well as for the reference set of all KEGG pathways, we calculated how many of them are significantly enriched within (i) T2D (GOLD) genes from the type 2 diabetes genetic association database (T2DGADB)(17) and (ii) a more recent and extended T2D GWAS dataset from a genome-wide meta-analysis (see Methods). We observed a significant enrichment of HiCc pathways in the two datasets. In particular, the fraction of enriched pathways in GOLD data (50 pathways overall) was significantly higher for HiCc pathways (24 out of 66 pathways) than for non-HiCc pathways (26 out of 120 pathways), with a two-tailed Fisher’s exact p-value of 0.038. Similarly, the fraction of enriched pathways in the GWAS dataset (49 pathways overall) was significantly higher for HiCc (28 out of 66 pathways) than for non-HiCc pathways (21 out of 120 pathways), with a two-tailed Fisher’s exact p-value of 0.00042 (Figure 3b). Overall, 55 HiCc pathways were enriched in T2D relevant omics data (Figure 3c). Among the pathways that were the most significantly enriched in the -omics data were several whose relevance to T2D is well established, such as Type II diabetes mellitus, PPAR signaling, insulin signaling, calcium signaling and chemokine signaling pathways (Supplementary Table 2). The role of chemokine signaling is known in T2D, as islet inflammation is involved in the regulation of β-cell function and survival in T2D (18). This indicates that our approach captures pathways that have T2D relevance in an unbiased way.

### Validation of HiCc pathway genes

Among the genes in the significant HiCc pathways, we found 51 eQTLs that pass the FDR<1% threshold and 10k permutation, as shown in Table 1, using an extended follow-up study to our original islet data, which consists of 89 pancreatic islet donors(19). In total, we observed a SNP within 250 kb up or downstream of the genes in 33 pathways. The enrichment of 33 pathways with the background distribution was significant (p-6.618e-13, odds ratio: 7.49), which indicates that we were able to capture the genomic signals among the HiCc pathways. We next tested the fold-change difference of eQTL genes vs. non-eQTL genes in the transcriptome data of rat islets pre-cultured with 2, 5, 10, and 30 mM glucose levels (GSE12817) (20). We found that the eQTL genes have significantly higher fold change compared to non-eQTL genes in 5 mM (Mann-Whitney test *p*=0.009), 10 mM (*p*=6.72e-05), and 30 mM (*p*=8.63e-06) glucose levels (Figure 4a). This indicates the potential role of eQTL genes in β-cell function as these cells are regulated both acutely and chronically by the extracellular glucose concentration. By applying a greedy algorithm (Steiner tree) in an integrated network of EGRN and protein interaction network, we observed a single connected component of eQTL genes with few linkers (grey nodes) (Figure 4b). This signifies that in reality a network environment is better characterized by the *local impact hypothesis (1)*, indicating that perturbations are localized to the immediate vicinity of the perturbed genes that carry the eQTLs.

**Figure 4:**
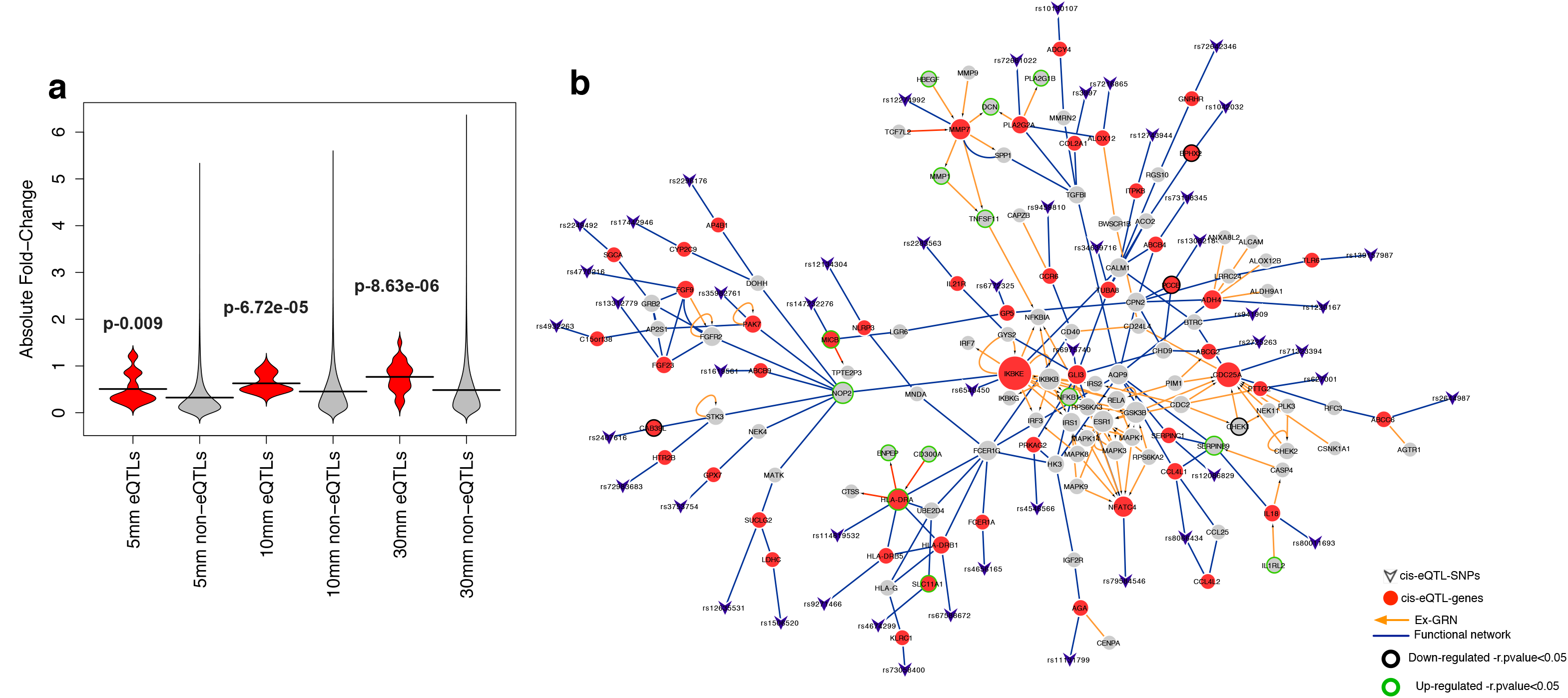
eQTLs and the functional network. a. eQTL genes and glucose levels: we tested the fold change difference of eQTL genes vs. non-eQTL genes in the transcriptomic data of rat islets pre-cultured at 2, 5, 10, and 30 mM glucose. eQTL genes are significantly changed in expression compared to non-eQTL genes. b. Integrating EGRN and gene interaction networks with the eQTL-gene relationship associated with T2D. Most of the genes in the integrated module are up-regulated in T2D subjects (nodes in green).

**Table 1:**
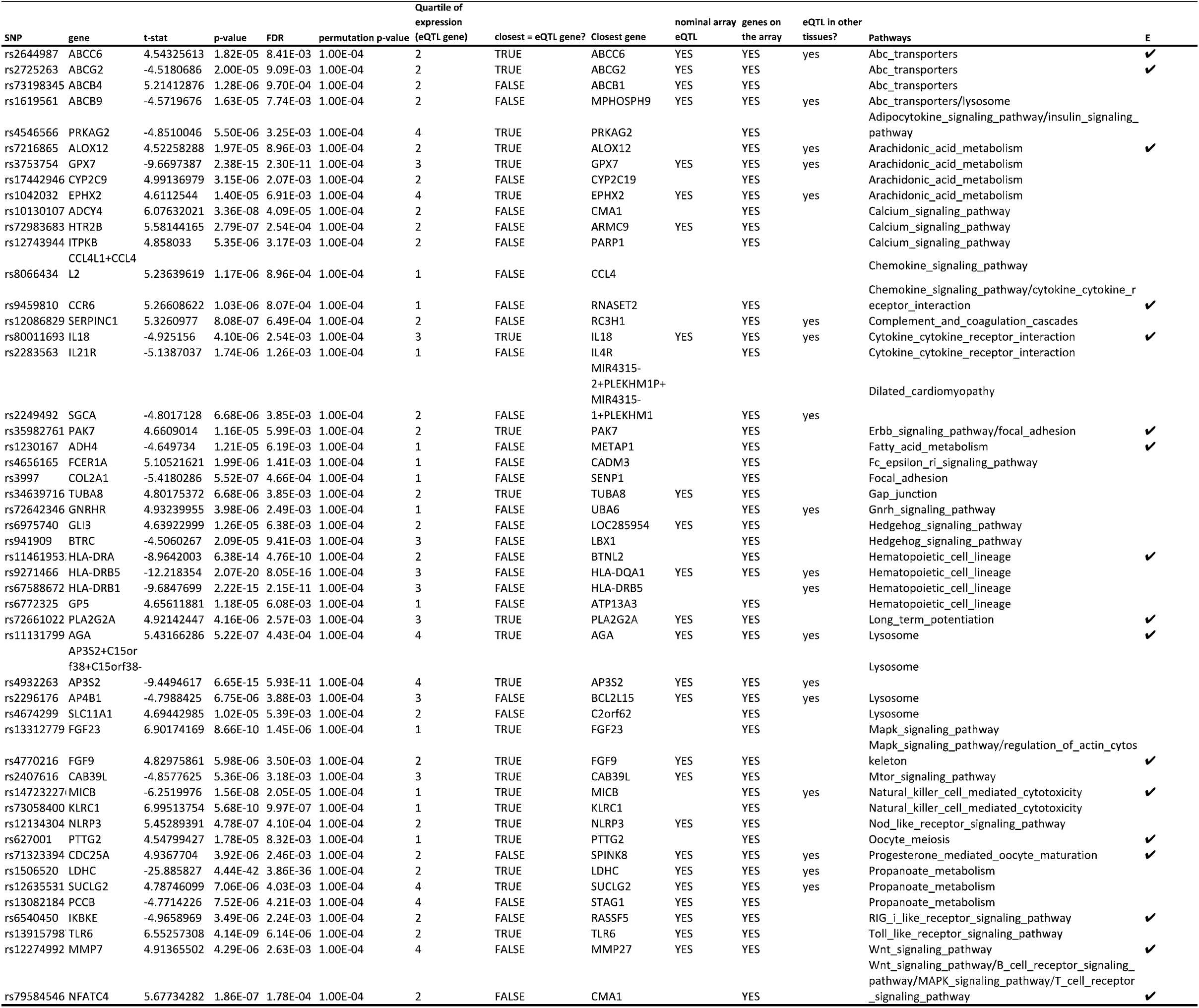
cis-eQTL in HiCc pathway genes

### New mechanistic connections in T2D

To find the connection between the HiCc pathways and T2D-associated genes, we focused on the eQTL gene (q-value=1.78E-04), NFATC4 (rs79584546), which is associated with four HiCc pathways: Wnt signaling, B-cell receptor signaling, MAPK signaling, and T-cell receptor signaling pathways (Figure 5a). We asked whether the T2D-associated genes from GWAS and literature are downstream targets of NFATC4. This might help in explaining their role in four HiCc pathways. The NFATC4 gene interacts with PPARG, and different MAP kinases (MAPK1, MAPK3, MAPK8, MAPK9, MAPK14) (Supplementary Figure 4). It is known that ablation of NFATC4 increases insulin sensitivity, in part, by sustained activation of the insulin-signaling pathway(21). We, therefore, next explored the downstream effect of transcriptional targets of NFATC4.

**Figure 5:**
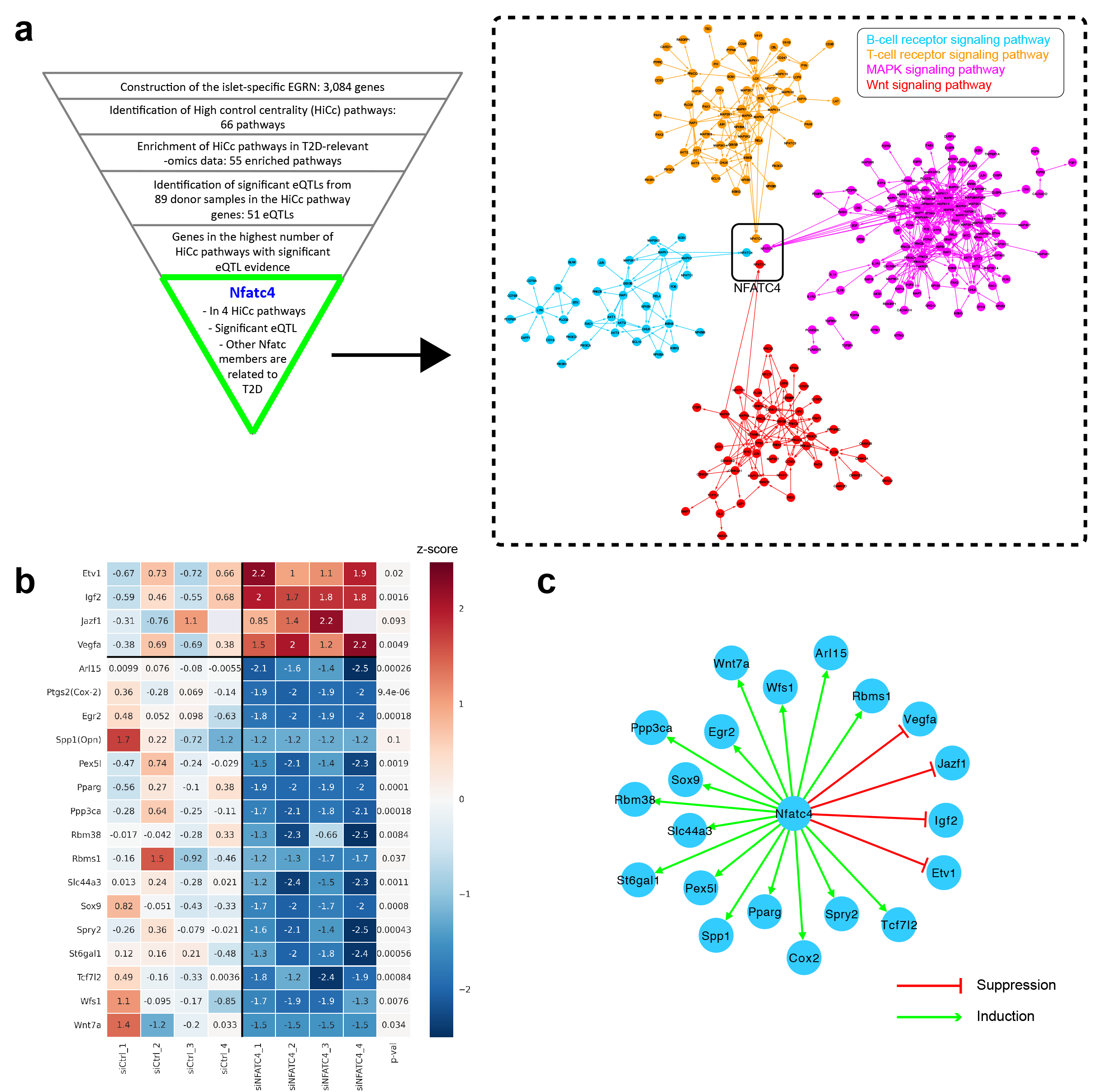
Nfatc4 *in vitro* validation. a. Nfatc4 is at the intersection of four HiCc pathways, namely B-cell receptor signaling pathway, T-cell receptor signaling pathway, MAPK signaling pathway, and Wnt signaling pathway. b. The effect of silencing of Nfatc4 on putative downstream genes. Colors indicate the Z-score, which was calculated across all samples per gene and is shown relative to the average Z-score of the control samples. P-values were obtained by a twosided t-test for two independent samples. c. The network of the putative downstream effect of Nfatc4 validated by *in vitro* silencing experiments.

NFATC4 has been reported to be a possible target for up-regulated transcription by TGF-alpha(22). Therefore, we used TGF-alpha to augment or induce Nfatc4 mRNA transcription during a glucose challenge in functional rat pancreatic beta islet cells *in vitro*. Nfatc4 was effectively silenced (see Methods for details on the *in vitro* silencing experiments) as seen in real time qPCR with 4 different rat-specific Nfatc4 probes (Supplementary Figure 5a-d). In order to assess the downstream effect of Nfatc4, we gathered putative T2D candidate genes regulated by the NFAT family members from GWAS and literature^23^. In particular, we selected 13 highly up- and down-regulated transcriptional targets of Nfatc1 and Nfatc2, namely Etv1, Jazf1, Pparg, Vegfa, Arl15, Pex51, Rbm38, Rbms1, Slc44a3, Spry2, St6gal1, Tcf7l2 and Wfs1, based on a recent report^23^ that they inhibit the expression of the first four genes while promoting the expression of the latter nine genes. We supplemented this list with seven putative transcriptional targets of Nfatc4, namely Cox2 (Ptgs2), Egr2, Igf2, Opn (Spp1), Ppp3ca, Sox9 and Wnt7a, through literature search via the MetaCore platform [https://portal.genego.com/]. The silencing of Nfatc4 in rat islet cell lines resulted in increased expression of, Etv1, Igf2, Jazf1 and Vegfa mRNA, compared to control siRNA treatment. In contrast, Arl15, Cox2, Egr2, Opn, Pexl5, Pparg, Ppp3ca, Rbm38, Rbms1, Slc44a3, Sox9, Spry2, St6gal1, Tcf7l2, Wfs1 and Wnt7a, all resulted in decreased mRNA expression post Nfatc4 silencing (Figure 5b).

The silencing of Nfatc4 in rat islet cell lines, thus displays reduced expression of 16 downstream T2D candidate genes and increased expression of 4 downstream T2D candidate genes. To complete the circle, we went back to the human islet data and assessed correlation of Nfatc4 expression with the aforementioned genes. We found that NFATC4 expression was positively correlated with ETV1, VEGFA, EGR2, RBM38, SOX9, ST6GAL1, TCF7L2, WFS1 and WNT7A and negatively correlated with SPRY2 (Supplementary Figure 6 and Supplementary Table 6). This indicated the potential influence of NFATC4 expression on the expression of the above genes in human pancreatic islets as well. NFATC4 expression also positively correlated with HbA1C levels, indicative of some effect on glycemic status. The mechanistic connection between NFATC4 and TCF7L2 is particularly of interest as TCF7L2 has been established as a major T2D susceptibility gene (23,24). TCF7L2 is also a member of two of our HiCc pathways, namely B-cell receptor signaling and Wnt signaling pathways. This indicates the possibility of finding further unexplored connections between the members of HiCc pathways within the context of specific diseases. Overall, the approach helps not only in identifying the potential dysregulated pathways, but also establishes the downstream regulation by NFATC4 in four important T2D pathways (Figure 5).

## Discussion

By exploiting the topological measures of cellular re-wiring associated with disease progression, it is possible to identify new disease genes and pathways (1,2,6). With the advances in control theory, and control principles becoming an important consideration in many disciplines, including disease biology and biological network analysis (25–27), network dynamics and regulation also provide opportunities to identify key regulatory genes in health and disease. Here, we exploited the *control centrality* measure to identify key pathways that could drive the islet regulatory network in T2D. We established a framework to find the pathways that might be related to the underlying hierarchical structure of disease regulation and is able to add a new dimension compared to the number of established methods like HotNet2(8). We identified 66 pathways as statistically significant in the analysis and were able to identify known T2D pathways (p=5.76E-05) among the top gene list in our analysis, which validates the approach as a means to capture disease relevant pathways. The pathways captured in our analysis were also enriched in T2D relevant omics sources. Furthermore, eQTL analysis helped in pruning the HiCc pathways genes by identifying the variants actually affecting the gene expression levels of these genes (i.e. cis-eQTLs). These eQTL genes showed glucose-induced changes in the islets.

Finally, we experimentally validated the process by which a particular eQTL gene (NFATC4) regulates the expression of numerous putative downstream T2D candidate genes of two other genes of the same family, NFATC1 and NFATC2, which were also shown to regulate T2D related genes by previous studies^23^. In particular, Nfatc4 silencing results confirmed similar transcription regulation pattern for these genes except for Igf2, which showed an opposite effect relative to the siControl condition. Results also demonstrated that Nfatc4 increases gene expression of Pparg, Tcf7l2, and Wfs1 which are genes already reported to be associated with type 2 diabetes as well as activation of the Wnt pathway as predicted by systems genetics approach that is subsequently validated *in vitro* (28). However, silencing Nfatc4 also appears to have a tendency to inhibit osteopontin (Opn). We have previously demonstrated that glucose dependent insulinotropic polypeptide (GIP) stimulates expression of OPN in human islets where OPN exerts protection against cytokine-induced apoptosis(29). The connection between NFATC4 and TCF7L2, which has not been reported previously in the literature, is particularly important as it adds to the mechanistic information on two pathways (Wnt signaling and B-cell signaling pathways) that were found to have high control centrality in the T2D pathobiologic context. Owing to this new connection, NFATC4 and TCF7L2 also emerge as potential players in the pathway communication between the T-cell receptor signaling, MAPK signaling, Wnt signaling, and B-cell signaling pathways. Overall, the positive experimental validation of our model shows the utility of the control centrality approach in pathway prioritization. In particular, it may pave the way for discovering hitherto uncovered cross-talk between pathways.

These results might help us understand better the controllability of complex networks and provide a basis for designing an efficient strategy for optimizing (normal) network control. There are, however, outstanding annotation and methodological challenges remaining, including low-resolution pathway-based knowledge, limited cell type-specific information, and incomplete annotation of next-generation pathways. Despite these hurdles, as the number and type of functional annotations increase, coupled with technological advances in analytical methods that provide better guidance for the utility of pathway analysis, confidence in the results will likely improve. Although the approach has been demonstrated using pancreatic islet gene-expression data, it can be used to interpret pathways for other complex diseases. Overall, controllability-based network analysis may be of broad use in dissecting complex diseases and in discovering novel therapeutics targets in this coming era of systems medicine.

## Methods

### Construction of gene regulatory network (GRN)

We constructed a disease gene-regulatory network (GRN) by integrating gene expression data from human pancreatic islets together with known information about transcription factor binding sites. We used gene expression data from islets from 63 cadaver donors provided by the Nordic Islet Transplantation Programme (http://www.nordicislets.org) (Supplementary Table 3). Islets were obtained from 54 nondiabetic donors (25 females, 29 males, age 59 ± 9, BMI 25.9 ± 3.5, HbA1c 5.5 ± 1.1) and 9 T2D donors (4 females, 5 males, age 57 ± 4, BMI 28.5 ± 4.5, HbA1c 7.2 ± 1.1). All procedures were approved by the ethics committee at Lund University. Purity of islets was assessed by dithizone staining, while measurement of DNA content and estimation of the contribution of exocrine and endocrine tissue were assessed as previously described (3). Gene expression was assayed using Affymetrix Human Gene 1.0 ST arrays. We normalized the data by robust multiarray averaging (RMA) and a custom Chip Description File (CDF) from the Michigan Microarray Lab http://brainarrav.mbni.med.umich.edu, Version 13) which helps estimate gene-level expression more accurately by summarizing probe sequences using up-to-date gene annotations.

To construct the GRN, we first used the LIMMA package in R(30) to select the most disease-relevant genes. For each gene, we computed the B-statistic for differential expression between the following phenotypic groups: 1) diabetic vs. non-diabetic, and 2) low vs. high levels of glycated hemoglobin (HbA1c). Expression analysis was carried out between 63 patients with and without T2D for (1) and donors with HbA1c < 6% and > 6% from the human islets mRNA data set for (2). A nominal *p*-value of < 0.05 was used to identify differentially expressed (DE) genes, yielding a set of 506 genes. We used a liberal threshold to obtain the signature genes that could be used for identifying the signature of disease as has been done previously (2). All genes were then ranked according to the average of the two B-statistics. We also added 48 genes that have been associated with T2D through GWAS (Supplementary Table 4) and that were also represented on the array to create a list of 554 total disease-relevant genes. Next, we computed the variance of the expression of each gene on the array across all 63 samples, and selected the top 2,000 most variable genes to add tissue-specific genes that are expressed and active in human islets.

The final list of 2,554 genes, together with the corresponding gene expression values over all 63 patients, was used as input to build the GRN. We applied Predictive Networks (PN) (31) a tool that gathers a comprehensive set of known gene interactions from a variety of publicly available sources as prior evidence, and then uses these known interactions together with gene expression data to infer gene regulatory networks via linear regression modeling(31). As prior evidence, we used known regulatory interactions from the TRANSFAC database and applied a weighting parameter of 0.75 to combine the regression and prior-based edges.

### Extended- gene-regulatory network (EGRN)

A gene can be involved in various interactions, and its role, and consequently its centrality, can vary across different biological networks. In order to obtain a higher resolution understanding of the signaling relationship in the GRN, we added kinase (http://www.phosphosite.org) and signaling events to the model (Signalink database) (32). This addition was done in order to add potential downstream signaling events by the nodes in the GRN. We call the full network, including transcriptional edges and signaling edges, the extended gene-regulatory network (EGRN).

### Ranking the KEGG pathways using the control centrality measure –HiCc pathways

We started with an un-weighted directed EGRN *G*=(*V, E*) with *N*=|*V*| nodes and *L*=|*E*| links. The *control centrality* of node *i*, denoted as *C*_c_(*i*), is defined to be the generic dimension of controllable subspace or the size of controllable subsystems if we control node *i* only (15). Hence, *C*_c_(*i*) captures the “power” of node *i* in controlling the whole network. For example, a simple network of *N*=7 nodes is shown in Fig.1B. When we control node *x*_1_ only, the controlled network is represented by a directed network with an input node *u*_1_ connected to *x*_1_. The dimension of the controllable subspace by controlling nodes *x_1_* is six, corresponding to the largest number of edges in all stem-cycle disjoint subgraphs (an example is shown in red in Figure 1), where “stem” is defined as a directed path starting from an input node, so that no nodes appear more than once in it. Hence, the control centrality of node *x*_1_ is *C*_c_(1)=6. In general, the control centrality of any node in a directed network can be calculated by solving a linear programming problem (15,33,34). The assumption was that if a pathway or module includes genes with high *C*_c_ values, it might be higher in the hierarchy and must regulate other downstream pathways with on average, lower *C*_c_ values. To test this hypothesis, for each pathway in the KEGG database, we identify the pathways with significantly greater *C*_c_ values compared to others. To calculate the statistical significance based on the *C*_c_ values of the genes representing the particular pathway, we use the Mann-Whitney U test with a cutoff *p*-value <0.05. A typical problem with pathway analysis methods is bias toward the enrichment of cancer-related pathways as these pathways have been studied more intensively. We, therefore, focus only on the non-cancer HiCc pathways.

### T2D relevance of the HiCc pathways in the EGRN

We next evaluated the relative enrichment of HiCc pathways genes across three T2D-specific datasets:

i. *Disease gene set*: T2D genetic association database (T2DGADB) aims to provide specialized information on the genetic risk factors involved in the development of T2D. 701 publications in the type 2 diabetes case-control genetic studies for the development of the disease were extracted (35), which was defined as the gold standard gene set. Overall, this dataset contained 143 genes.
ii. *Genomics*: This genome-wide meta-analysis (“DIAGRAMv3”) includes data from 12,171 cases and 56,862 controls of European descent imputed up to 2.5 million autosomal SNPs(36). We computed a single p-value for each gene in the interactome by the VEGAS method using the whole GWAS data set (37). We considered the genes with uncorrected p-value < 0.01 in our analysis, resulting in 1308 genes. There was little overlap between (i) and (ii) gene sets. Moreover, to avoid circularity, we exclude the 48 GWAS genes used in the construction of EGRN from this dataset.

The enrichment of HiCc, non-HiCc and all pathway genes with the above gene sets was calculated through Fisher’s Exact test.

### eQTL analysis of HiCc pathway genes

One of the major findings from the T2D GWAS is that most of the trait-associated SNPs are located in intronic, intergenic, or other non-coding regions of the genome (38). As many SNPs are located in noncoding regions, suggesting they may influence gene expression, we analyzed whether any SNP within 250 kb of HiCc pathway genes (cis) would influence their gene expression (eQTL). We used a linear model adjusting for age and sex as implemented in the R Matrix eQTL package. Genotyping was performed on the Illumina HumanOmniExpress 12v1C chips, and all the samples passed standard genotype QC metrics. Genotypes were imputed to 1000 Genomes data, using IMPUTE2 and SHAPEIT.

The transcriptomic data from rat pancreatic islet after culture in low, intermediate and high glucose was retrieved from Gene Expression Omnibus (GEO-GSE12817). We performed differential expression analysis between 2 and 10, 2 and 30, 5 and 10, 5 and 30 or 10 and 30 mmol/l glucose. We used the limma R package (ver 3.10.1) for the differential expression analysis. We compared the fold change (absolute log) of eQTL genes to all other differentially expressed genes and computed the p-values by applying Mann-Whitney U test.

### Functional network and eQTL-HiCc pathway genes

To evaluate the impact of genes in the vicinity of eQTL in different sources of network data, we used HumanNet gene-interaction data. HumanNet uses a Näive Bayesian approach to weight different types of evidence together into a single interaction score focusing on data collected in humans, yeast, worms, and flies(39)._ The hypothesis we tested was that if the gene products (e.g., proteins) linked to the same disease phenotype interact with each other more often than randomly linked gene products (40–42) and cluster in the same network neighborhood, then eQTL genes must be connected through a single component in the gene interaction network. We applied a heuristic Steiner tree algorithm to find the minimum number of genes that can connect the eQTL into a connected component in the gene-interaction network.

### Nfatc4 silencing and stimulation experiments *in vitro*

To validate Nfatc4 effect on putative downstream T2D candidate genes, Nfatc4 mRNA expression was silenced in clonally derived rat pancreatic β-islet cell line INS-1 (832/13), a generous contribution from Dr. Rohit Kulkarni. Cells were maintained on RPMI 1640, 10% fetal calf serum, 10 mM HEPES, 2 mM L-glutamine, 1 mM sodium pyruvate and 0.05 mM 2-mercaptoethanol (Thermo Fisher) supplemented with penicillin (100 Units/ml) and streptomycin (100 μg/ml) (Pen/Strep). We transfected rat specific Nfatc4 siRNA (Rat Nfatc4 ON-TARGET Smart Pool siRNA, GE Dharmacon) using Lipofectamine RNAiMAX (Thermo Fisher) as per manufacturer’s recommendation. For controls, we employed ON-TARGETplus Non-Targeting Control Pool (GE Dharmacon) using the same concentration as the test siRNA. Silencing was carried out in each well containing 60% confluent cells in culture and were exposed to transfecting medium for 48 hours at 37°C in a humidified atmosphere containing 95% air and 5% CO2 in the presence of 20 pmol siRNA per well in 12-well cell culture microplates.

After silencing, cells were allowed to recover for 16 hours in regular growth media followed by stimulation of Nfatc4 signaling by TGF alpha (TGF-a) at 50 ng/ml as described in another publication (22) for 6 hours. Subsequently, cells were washed with HBSS once, then exposed to 17.3 mM glucose in HBSS for 1.5 hours. RNA from cells were harvested using column isolation (GE Healthcare Life Sciences). Putative genes downstream of Nfatc4 were tested using probes from Taqman Gene Expression Assay system (Thermo Fisher) listed in Supplementary Table 5. Real time PCR was done with Biomark HD (Fluidigm) thermocycler. Quadruplicates were used per condition.

### Expression correlation of NFATC4 with putative downstream genes in human islet cells

RNA sequencing was performed on the Hi-seq as described previously (19). Alignments were performed using STAR and gene counts were assessed using feature counts. Spearman correlation was used to assess the relationships between NFATC4 expression and ETV1, IGF2, JAZF1, VEGFA, PTGS2/COX2, EGR2, SPP1 (OPN), PPARG, PPP3CA, RBM38, RBMS1, SOX9, SPRY2, ST6GAL1, TCF7L2, WFS1 and WNT7A.

## Acknowledgments

This work was supported by National Institutes of Health (NIH) grants P50-HG004233-CEGS, MapGen grant (U01HL108630) and P01 HL083069, U01 HL065899, P01 HL105339, R01HL111759, and RC HL10154301.

## Author contributions

A.S, A.H. and J.M. conceived the idea and designed the experiments; A.S., A.H., J.M., M.S., Y.Y.L., J.F., R.B.P and M.S. performed the experiments and analysis. M.P. constructed the gene regulatory network. J.L.D. designed and performed the Nfatc4 in vitro knockdown experiments. A.S., A.H., A.L.B., S.T.W., E.K.S., M.A, L.G, M.V., J.L. wrote and modified the manuscript.

